# Tissue destruction during food spoilage is associated with the formation of biofilms by *Pseudomonas* species

**DOI:** 10.1101/2025.07.11.664243

**Authors:** Laura M. Nolan, James J. Lazenby, Ryan Sweet, Gregory J. Wickham, Haider Al-Khanaq, Samuel J. Bloomfield, Alison. E. Mather, Cynthia B. Whitchurch

## Abstract

Members of the *Pseudomonas* genus are common spoilers of a range of meat, dairy and vegetable products. While we have a good understanding of the *Pseudomonas* species typically responsible for spoilage, we know very little about how these bacteria interact with food surfaces during spoilage. Here we assessed the spoilage capabilities of a large panel (n=124) of *Pseudomonas* species food-derived isolates on meat (chicken) and leafy greens (spinach). Most isolates (71/124) were capable of spoiling both foods, but some were only capable of spoiling only chicken (21/124) or spinach (23/124), or neither (9/124). Our data also demonstrated that the type of fresh food the strain was isolated from influenced spoilage capabilities: strains isolated from meat were likely to spoil both chicken and spinach, while those isolated from leafy greens were more likely to spoil only spinach. We used fluorescence microscopy to visualise how *Pseudomonas* spoilage species interacted with the meat or leaf tissue and observed significant tissue destruction associated with biofilm formation. For chicken, this was associated with the formation of dense biofilm pillars that penetrated deep into the tissue. For spinach we observed biofilms on the leaf in areas of tissue degradation. Finally, we explored the correlation between potentially relevant phenotypes (*in vitro* biofilm, motility and secreted enzyme production) and spoilage capabilities. There was no significant correlation between any of these phenotypes and spoilage, except for secreted protease activity and chicken spoilage. Overall, this study increases our understanding of processes involved in food spoilage by *Pseudomonas* species.

## Introduction

Almost 10% of the world’s population do not have enough to eat (FAO, 2020). Given that the world’s population is predicted to reach 9.8 billion by 2050, more than a 50% increase in food availability is needed to feed the growing population (WRI, 2018).

Despite these drastic numbers approximately 17% of food produced for human consumption worldwide is lost or wasted (UNEP, 2021). Food loss is defined as food products being discarded prior to the retail stage, for instance during production, harvesting, and preparation, whilst food waste is defined as food products being discarded at the retail, food service provider, consumer and household stages (World Food Program USA (WFP USA), 2022). These processes also squander resources such as water, energy and land that go into producing food. Reducing food loss and waste is one of the Sustainable Development Goals set by the United Nations, which aims to significantly increase global sustainability by 2030 (United Nations, 2021).

Food spoilage is a major contributor to food loss and waste, and is due to physical, chemical and/or microbial factors that cause changes in the texture, smell, taste or appearance of food, leading to it be discarded (Mafe et al., 2024). Many foods provide an ideal environment for growth and proliferation of certain microbes. During growth, these microbes can produce a range of molecules that cause food spoilage, including by-products of metabolism that produce off-odours, off-flavours and discolouration; production of extracellular polymeric substances that cause a slimy texture; and secretion of enzymes that break down food, causing malodour and changes in texture and flavour (Alegbeleye et al., 2022; Barth et al., 2009; Braun et al., 1999).

Non-pathogenic *Pseudomonas* species, like *P. fluorescens, P. fragi* and *P. putida*, are common spoilers of a range of meat, poultry, seafood, vegetable, fruit and dairy products (Karanth et al., 2023). These species are capable of growing at a wide range of temperatures, including cold storage temperatures (Raposo et al., 2016).

Food spoilage by these species occurs due to the production of secreted enzymes, such as proteases, lipases and cellulases; production of extracellular polymeric substances such as proteins, extracellular DNA (eDNA) and polysaccharides, which creates a slime layer on food; production of pigments, such as pyoverdine (yellow-green) and pyomelanin (brown-black), which result in food discolouration; and production of volatile organic compounds (VOCs), which can result in off-flavour and off-odour of foods (Karanth et al., 2023; Kumar et al., 2019; Wickramasinghe et al., 2020).

Food spoilage microbes grow on food in complex 3D communities called biofilms (Korber et al., 2009). A biofilm is a community of microbes that are encased within an extracellular matrix/slime, composed of exopolymeric substances (EPS), that attaches microbial cells to a surface and/or to one another (Sauer et al., 2022).

Biofilm development typically begins with attachment of microbes to a biotic or abiotic surface (Stoodley et al., 2002). During biofilm development microbes produce EPS that facilitate cell-cell and cell-substratum interactions within the biofilm matrix, to grow the structure and maintain integrity. The most common EPS matrix components are polysaccharides, extracellular DNA (eDNA) and proteins (Flemming et al., 2016).

Biofilms are a protected mode of growth that allows microbes to thrive in hostile environments, by providing protection from antibiotics, disinfectants and immune system factors (Flemming et al., 2016). Consequently, biofilms are extremely difficult to eradicate once they have formed (Singh et al., 2017). Given this, significant efforts have been made to prevent biofilm formation, or to disrupt biofilms once they have formed in a range of settings, including the food chain (Galié et al., 2018). In this setting, a major focus has been on targeting biofilms on food processing surfaces using chemical and enzymatic treatments, as well as the use of microbial-derived antibacterial compounds (bacteriocins, such as nisin, and biosurfactants) or essential oils obtained from plants (Galié et al., 2018).

Despite significant efforts to target biofilms in the food chain, food spoilage remains a major global challenge. One contributing factor is that while biofilms have been well-studied under *in vitro* conditions, and on abiotic food chain surfaces like stainless steel and plastic (Carrascosa et al., 2021), biofilms on the surface of food have not been extensively studied. To our knowledge no studies have characterised *Pseudomonas* spoilage species biofilms on post-harvest leafy greens. Two prior studies have visualised *Pseudomonas* spoilage species biofilms on the surface of beef or pork, from when the meat was fresh until spoiled (Delaquis et al., 1992; Wickramasinghe et al., 2019). These studies mainly focused on visualising biofilm structures on the meat surface, i.e. above the tissue and reported a range of distinct morphological characteristics for different *Pseudomonas* species over the course of spoilage.

In this study we characterised spoilage capabilities of a large panel (n=124) of *Pseudomonas* species food-derived isolates on fresh skinless chicken breast and fresh baby spinach. Given that spoilage is associated with significant alterations in the meat or leaf tissue we wanted to understand how *Pseudomonas* spoilage species interacted with the respective tissue during spoilage. We found that biofilm formation was associated with significant destruction of both tissue types and in the case of chicken, with deep penetration into the tissue. We also explored the correlation between potentially relevant phenotypes (*in vitro* biofilm, motility and secreted enzyme production) and spoilage capabilities. There was no correlation between any of these phenotypes and spoilage of chicken or spinach, except for secreted protease activity and chicken spoilage. Overall, this study increases our understanding of how *Pseudomonas* spoilage species interact with fresh meat and leafy greens to cause spoilage and will be used as a foundation for development of interventions that extend food shelf life and reduce food waste.

## Materials and Methods

### Bacterial strains and culture conditions

The *Pseudomonas* strains used in this study were isolated from fresh meat, poultry, seafood, and leafy green vegetable samples that were sampled as part of a food study (Janecko et al., 2021). Isolation of *Pseudomonas* species and species/strain determination was carried out in a follow up study (Bloomfield et al., 2024) or in the current study. For the latter, isolation and species/strain identification was performed as described in Bloomfield et al., 2024. See Supplementary Table 1 for the full list of isolates. *Pseudomonas* isolates were cryogenically stored at-70°C in 15% (v/v) glycerol, cultured on lysogeny broth (LB) solidified with agar at 1.5% (w/v) for routine maintenance and grown in LB prior to use in assays where specific media was used, as detailed below.

### Food spoilage assays

Skinless chicken breast and baby spinach (*Spinacia oleracea*) leaves were obtained from the a supermarket in Norwich, United Kingdom, on the day of the experiment. A cold storage chain was maintained from the supermarket to the lab. To remove the native microbiota from the food surface, the chicken or spinach leaves were washed in sodium hypochlorite (0.1%, [v/v]) followed by a wash in sterile deionised water, with each wash repeated twice. Each food product was then exposed to UV in a Stratalinker UV Crosslinker 2400 machine (Stratagene) for a period of 10 min (chicken) or 5 min (spinach) at 3000 µjoules. For chicken the bleached outer surface was removed with a scalpel aseptically, and the sterile inner tissue was cut into small strips which were subjected to additional UV treatments (on each side), then further cut with a 8 mm biopsy punch to generate pieces of chicken that were approximately 8 mm in diameter and 6 mm high, which was subjected to a final UV treatment on each side before use. For spinach only the leaf region (i.e. not stem or rib region) was used and was cut into pieces that were approximately 6 mm in diameter, which was UV treated on each side prior to use. The sterilised food samples were inoculated with a *Pseudomonas* isolate as follows. Each isolate was grown in LB at 25 °C, shaking at 200 rpm, overnight. Prior to inoculation the overnight culture was diluted 1:100 into Phosphate Buffered Saline (PBS) and one piece of sterile chicken or spinach added to the cell suspension. This was incubated at room temperature (approx. 22 °C) for 1 h after which the cell suspension was then removed. Sterilised chicken and spinach were also inoculated with PBS in the absence of bacterial cells as controls. These chicken and spinach samples were incubated at room temperature (∼ 22 °C) under humid conditions by placing the samples in a plastic box which contained paper towel dampened with 10 mM Mg_2_SO_4_. Spoilage scored at 2 and 7 days. A Sensory Evaluation assessment was used by a panel of trained individuals to score changes in the smell, colour and texture of the chicken and spinach (de Bouillé and Beeren, 2016) compared to the uninoculated control.

Spoilage was recorded as unspoiled if it was identical to the uninoculated, sterilised control (-), or assessed as a low (+), medium (++) or high level (+++) of spoilage.

### Imaging food spoilage biofilms

A stereomicroscope (M165c; Leica) with a colour CCD camera (DFC450; Leica) and LAS-X software (Leica) was used to visualise and record spoilage of chicken and spinach samples that had not been stained with any dyes. To visualise biofilms formed by live cells, the tetrazolium redox-based dyes, 5-Cyano-2,3-Ditolyl-Tetrazolium Chloride (CTC; Merck) and 2,3,5-triphenyl-2H-tetrazolium chloride (TTC; Merck) were used to visualise biofilms on chicken and spinach, respectively.

Respiring cells reduce these dyes to CTC-or TTC-formazan crystal, which is visualised in the red fluorescent channel or as red visible light, respectively. To stain biofilms, CTC (1.5 mg/mL [w/v] in PBS) or TTC (0.5% [w/v] in PBS) was added to the chicken or spinach sample, respectively. The samples were fully immersed in the dye solution and incubated at room temperature for 1 h, protected from light. For chicken, the samples were also stained with 4’,6-diamidino-2-phenylindole (DAPI) (2 µg/mL [w/v], Merck) and/or ethidium homodimer-III (EthHD-III) (5 µM, Biotium) for 30 min at room temperature, protected from light. The dyes were then removed and the sample washed twice with PBS. The sample was then fixed with 4% (w/v) paraformaldehyde (PFA) overnight at 4 °C. The next day the PFA was removed, and the sample was washed twice with PBS prior to imaging.

For chicken, the sample was then transferred to a μ-Plate 24 well glass bottom plate (Ibidi) and imaged with fluorescence microscopy on a DeltaVision Elite Widefield inverted microscope (Image Solutions) with a PCO Edge 5.5 monochrome sCMOS camera (Excelitas) and a Spectra 7 Light Engine (Lumencor) using a x10 UPlanFLN Semi Apochromat objective (Olympus), SoftWoRx acquisition software (Applied Precision), and filter sets for DAPI (excitation 395 nm/25 nm, emission 455 nm/50 nm), EthHD-III (excitation 550/15, emission 605/52) and CTC (excitation 470 nm/24 nm, emission 605 nm/52 nm). Deconvolution of the widefield image was performed in SoftWoRx using an enhanced additive deconvolution method with 20 iterations.

These widefield deconvolution images were converted to either 2D maximum intensity projections (MIPs) using Fiji 2.16.0/1.54p software (Schindelin et al., 2012) or 3D images using Imaris 10.2.0 software (Bitplane) and the normal shading 3D volume rendering mode. For spinach, the sample was then transferred to a 24 well glass bottom µ-Plate and imaged using light microscopy on an Olympus IX71 microscope with a colour CMOS camera (Elite-5 Cytocam, MicroPix) and x10 UPlan FL N objective (Olympus). Fiji (Schindelin et al., 2012) was used to generate images for publication.

### *In vitro* biofilm assay

Overnight cultures grown in LB at 25 °C and shaking at 200 rpm were used for microtitre plate crystal violet biofilm assays. Overnight cultures were diluted 1:100 into 1x M63 media (3 g/L KH_2_PO_4_, 7 g/L K_2_HPO_4_ and 2 g/L (NH_4_)_2_SO_4_) supplemented with MgSO_4_.7H_2_O (1 mM), glucose (0.2% [w/v]) and casamino acids (0.5% [w/v]) into a well of a 96-well microtitre plate (Nunc) and incubated statically at 25 °C for 24 h. The microtitre plate lid was replaced with a breathable Aeraseal membrane. The biofilm biomass attached to the microtitre plate well was stained with crystal violet (0.1% [v/v]), and extracted using acetic acid (30% [v/v]) with the absorbance (A595 nm) of the extracted crystal violet allowing quantification of biofilm biomass, as described previously (O’Toole, 2011).

### Secreted protease assay

Overnight cultures grown in LB at 25 °C and shaking at 200 rpm were used for secreted protease skim milk agar assays, as described previously (Brown and Foster, 1970), with some modifications. For each strain 5 µL of overnight culture was spotted onto the surface of a skim milk agar plate (20 mL total volume/plate). The skim milk agar plates were composed of 1x M63 (as above) supplemented with MgSO_4_.7H_2_O (1 mM) and ultra-high temperature (UHT) skim milk (15% [v/v]) and solidified with agar (1.5% [w/v]). Each plate was incubated at 25 °C for 48 h. The longest (a) and shortest (b) diameters of each cleared zone were measured and the average diameter used to calculate the surface area using the formula: area=π(d/2)^2^.

### Secreted lipase assay

Overnight cultures grown in LB at 25 °C and shaking at 200 rpm were used for secreted lipase assays. Cells were pelleted by centrifugation (12,000 *g*, 5 min, room temperature) and 10 µL of supernatant used in a secreted lipase assay, as described previously (Wretlind et al., 1977). Briefly, *p*-nitrophenyl caprylate (7 mM) was used as a substrate. 1 µL of this substrate was mixed with 100 µL of NaPO_4_ buffer (0.1 M, pH 7.4) and added to the culture supernatant in a 96 well microtitre plate and incubated at 25 °C in a plate reader (Tecan). Secreted enzyme activity was determined from the linear increase in A410 nm was measured over time for that period at 25 °C. The extinction coefficient (18,000 M^-1^) for the product, *p*-nitrophenol, the pathlength (0.28 cm) and reaction volume were used to convert the slope in absorbance units/min to moles/min.

### Secreted cellulase assay

Overnight cultures grown in LB at 25 °C and shaking at 200 rpm were used for secreted cellulase assays. For each strain 5 µl of overnight culture was spotted onto the surface of a carboxymethyl cellulose (CMC) agar plate (20 mL total volume/plate). These plates were composed of NaNO_3_ (0.05% [w/v]), K_2_HPO_4_ (0.05% [w/v]), KCl (0.05% [w/v]), MgSO_4_.7H_2_O (0.025% [w/v]), yeast extract (0.025% [w/v]), glucose (0.05% [w/v]), agar (1% [w/v]), CMC (1% [w/v]) and Congo Red (80 μg/mL in dH_2_O). The plates were incubated at 25 °C for 48 h. After incubation the plates were flooded with dH_2_O and the top colony removed with a sterile loop to reveal any clearing zone in the plate that did not extend beyond this zone. The longest (a) and shortest (b) diameters of each cleared zone were measured and the average diameter was used to calculate the surface area using the formula: area=π(d/2)^2^.

### Twitching motility assay

Interstitial biofilm expansion was assayed using a modification of the subsurface twitching motility stab assay described previously (Semmler et al., 1999). Briefly, the *Pseudomonas* strain to be tested was stab inoculated through an agar plate and cultured for 25 °C for 48 h. After incubation the longest (a) and shortest (b) diameters of each interstitial biofilm at the agar and petri dish interface were measured and the average diameter used to calculate the surface area using the formula: area=π(d/2)^2^.

### Swimming motility assay

Overnight cultures grown in LB at 25 °C and shaking at 200 rpm were used for swimming motility assays. For each strain 2.5 µL of overnight culture was spotted onto the surface of a swimming motility agar plate (15 mL total volume/plate). These plates were composed of yeast extract (0.5% [w/v]), tryptone (1% [w/v]), NaCl (0.5% [w/v]) and agar (0.3% [w/v]). Plates were incubated at 25 °C for 16 h. After incubation the longest (a) and shortest (b) diameters of each swimming zone were measured and the average diameter used to calculate the surface area using the formula: area=π(d/2)^2^.

### Bioinformatics

We performed tBLASTn analyses using translated *Pseudomonas* species nucleotide gene sequences (accession numbers in parentheses): *aprX* (PQ442365.1); *aprA* (NC_002516.2:1355631-1357070); *piv* (NC_002516.2:4671319-4672707); *lasA* (NC_002516.2:2032695-2033951); *lasB* (NC_002516.2:c4170483-4168987); *paaP* (NC_002516.2:3294282-3295892), *tliA* (AF083061.1), *lipA2* (NZ_LT907842.1:c1338025-1337378); *lipC* (FM163375.1) and *bcsZ* (NZ_CP169744.1) with the whole genome sequences for all isolates in our panel. A 70% amino acid identity and 70% coverage cut off was used to score for gene presence/absence.

### Statistical tests

All analyses were performed using GraphPad Prism v.10.4.1. Multiple logistic regression analyses for spoilage capabilities and *in vitro* phenotypic data had chicken or spinach spoilage outcome (i.e. spoilage or no spoilage) as the dependent variable and *in vitro* assay data (biofilm, secreted enzyme assays (protease, lipase and cellulase), twitching motility or swimming motility) as the independent variables (Table 1). Multiple logistic regression analyses for secreted enzyme gene presence/absence and *in vitro* phenotypic data had gene presence/absence as the dependent variable and *in vitro* secreted enzyme assay data (protease, lipase and cellulase) as the independent variables (Supplementary Table 4). Pearson’s correlation coefficient (r) analyses had the binary variables gene presence/absence and chicken or spinach spoilage outcome (i.e. spoilage or no spoilage) (Supplementary Table 5).

**Table 1.** Multiple Logisitic Regression Analyses.

| *Independent variable (n=124)* | *Coefficient* | *Standard error* | *95% CI* | *Z value* | *p value* | *Odds ratio* | *95% CI* |
| --- | --- | --- | --- | --- | --- | --- | --- |
| *In vitro* biofilm | -0.01844 | 0.1675 | -0.3400 to 0.3504 | 0.1101 | 0.9123 | 0.9817 | 0.7117 to 1.420 |
| **Secreted protease** | 0.01877 | 0.005668 | 0.008940 to 0.03157 | **3.311** | **<0.001** | **1.019** | **1.009 to 1.032** |
| Secreted lipase | 0.06281 | 0.04152 | -0.005086 to 0.1635 | 1.513 | 0.1303 | 1.065 | 0.9949 to 1.178 |
| Twitching motility | 0.001541 | 0.004861 | -0.002269 to 0.03646 | 0.3171 | 0.7512 | 1.002 | 0.9977 to 1.037 |
| Swimming motility | -0.0009832 | 0.0007197 | -0.002464 to 0.0004002 | 1.366 | 0.1719 | 0.999 | 0.9975 to 1.000 |
| *Hypothesis tests* | *Statistic* | *p value* |  |  |  |  |  |
| Log likelihood ratio (G squared) | 35.69 | <0.0001 (****) |  |  |  |  |  |
| *Pseudo R squared* |  |  |  |  |  |  |  |
| Tjur's | 0.257 |  |  |  |  |  |  |

**Spinach spoilage**
| *Independent variable (n=124)* | *Coefficient* | *Standard error* | *95% CI* | *Z value* | *p value* | *Odds ratio* | *95% CI* |
| --- | --- | --- | --- | --- | --- | --- | --- |
| *In vitro* biofilm | 0.007714 | 0.1558 | -0.2955 to 0.3496 | 0.0495 | 0.9605 | 1.008 | 0.7442 to 1.418 |
| Secreted protease | 0.007211 | 0.003749 | 0.0004496 to 0.01526 | 1.923 | 0.0544 | 1.007 | 1.000 to 1.015 |
| Secreted lipase | 0.05604 | 0.03344 | -0.001821 to 0.1326 | 1.676 | 0.0938 | 1.058 | 0.9982 to 1.142 |
| Secreted cellulase | 0.0092 | 0.01001 | -0.007342 to 0.03401 | 0.9186 | 0.3583 | 1.009 | 0.9927 to 1.035 |
| Twitching motility | 0.001174 | 0.002043 | -0.001292 to 0.01182 | 0.5747 | 0.5655 | 1.001 | 0.9987 to 1.012 |
| Swimming motility | -0.0006484 | 0.0007434 | -0.002138 to 0.0008342 | 0.8721 | 0.3831 | 0.9994 | 0.9979 to 1.001 |
| *Hypothesis tests* | *Statistic* | *p value* |  |  |  |  |  |
| Log likelihood ratio (G squared) | 16.93 | <0.01 (**) |  |  |  |  |  |
| *Pseudo R squared* |  |  |  |  |  |  |  |
| Tjur's | 0.1172 |  |  |  |  |  |  |

## Results

The strains used in this study consisted of 124 *Pseudomonas* strains that had been isolated from fresh meat, seafood and leafy greens (Supplementary Table 1). This panel consisted of *P. fluorescens* (54 isolates), *P. putida* (23 isolates), *P. fragi* (20 isolates) and *P. aeruginosa* (14 isolates), with fewer isolates of *P. koreensis* (5 isolates), *P. furukawaii* (3 isolates), *P. veronii* (2 isolates), *P. trivialis* (1 isolate), *P. alkylphenolica* (1 isolate) and *P. chlororaphis* (1 isolate). We assessed the spoilage capabilities of this panel of *Pseudomonas* isolates by inoculating each isolate onto fresh, in-date skinless chicken breast and baby spinach that had been sterilised (i.e. lacking the native microbiota) and incubated under humid conditions for 2 or 7 days at room temperature (∼22 °C). Uninoculated sterilised chicken and spinach was used as a control, which showed no signs of dessication or other alterations in the tissue for the duration of the experiment. At each time point spoilage was scored using a Sensory Evaluation assessment with the categories: unspoiled if it was identical to the uninoculated, sterilised control (-), spoiled to a low (+), medium (++) or high level (+++) (Figure 1; Supplementary Table 2).

**Figure 1.**
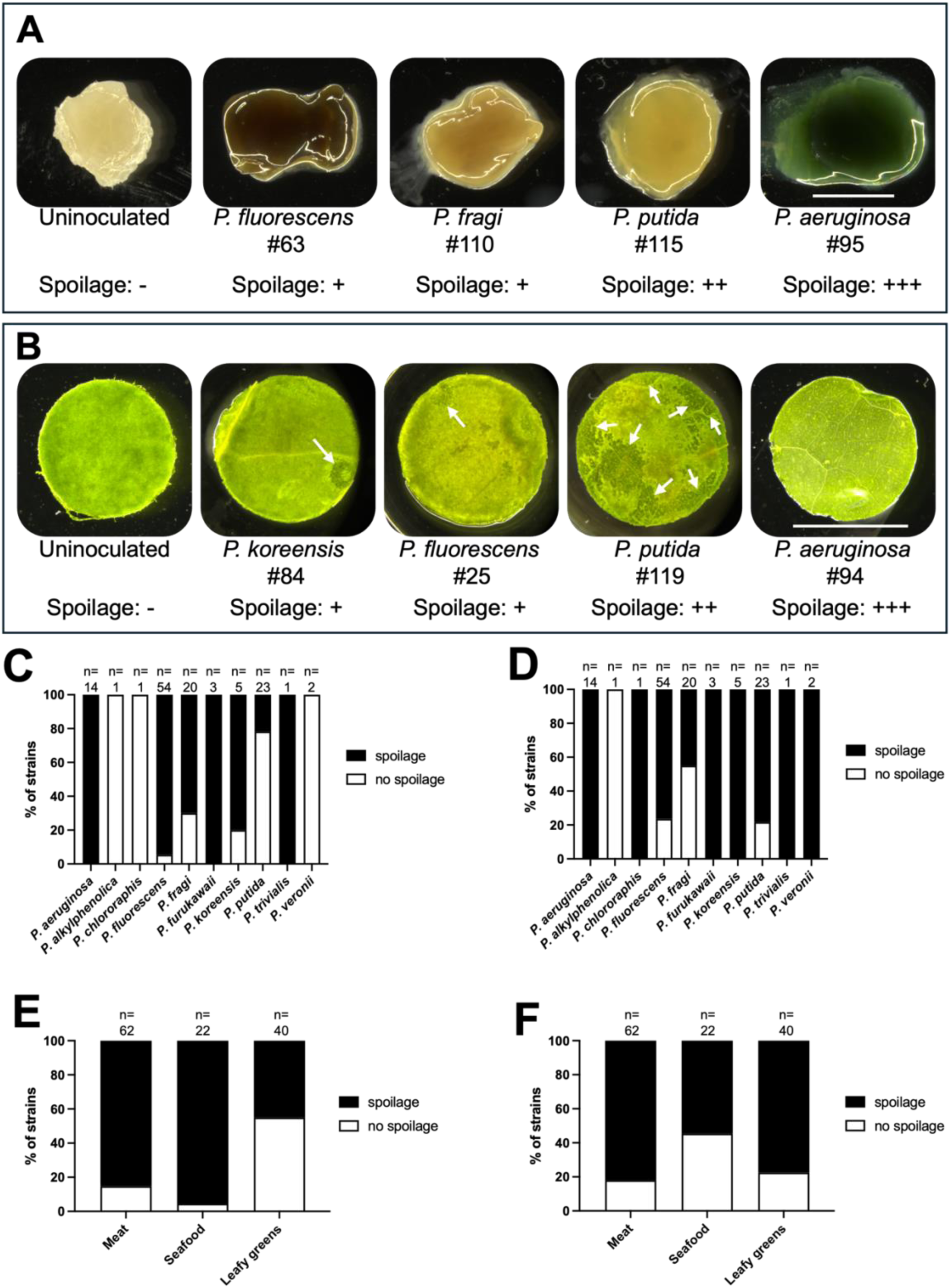
*Pseudomonas* isolate spoilage capabilities. (A-B) Stereomicroscope images of spoilage caused by select *Pseudomonas* food isolates on (A) chicken or (B) spinach. (C-D) Spoilage capabilities of *Pseudomonas* isolates for (C) chicken or (D) spinach. (E-F) Spoilage capabilities according to the type of food the strain was isolated from for (E) chicken or (F) spinach. For all samples, chicken and spinach was sterilised to remove the native microbiota and a single *Pseudomonas* isolate inoculated onto the food. The samples were incubated at ∼22 °C for 2 or 7 days and spoilage capabilities scored with (-) for no spoilage; and (+), (++) or (+++) for low, moderate and high degrees of spoilage, respectively. The full dataset is in Supplementary Table 2. For A-B: scale bar = 5 mm. In B white arrows indicate regions where there has been degradation of the cuticle and upper epidermis to reveal the mesophyll layer. NB: the entire cuticle and upper epidermis were completely degraded in highly spoiled samples (e.g. strain #94; +++). For C-F the number above each bar is the number of isolates from the species (C-D) or food type (E-F).

When chicken spoilage occurred, we saw production of slime and pigments (Figure 1A). We also observed loss of tissue structure in samples that were moderately spoiled or above (scored as ++ or +++) and complete loss of integrity and gelatinous texture in highly spoiled samples (scored as +++) (Figure 1A; Supplementary Table 2). In some cases, we also saw pigment production in conjunction with spoilage, e.g. dark brown pigment (Figure 1A, strain #63), which is likely to be pyomelanin (Fava et al., 1993); or blue/green pigment (Figure 1A, strain #95), which is likely to be pyocyanin (Stanley, 1947). For the uninoculated control we saw no pigment or slime production and no changes in tissue integrity (Figure 1A), demonstrating that these changes in the spoiled samples were due to microbial activity.

When spinach spoilage occurred we saw varying degrees of degradation of the cuticle and upper epidermis, which was scored as low or medium levels of spoilage (scored as + or ++), and where there was complete degradation of these layers to reveal the leaf mesophyll this was scored as high level spoilage (scored as +++), (Figure 1B; Supplementary Table 2). For the uninoculated control, the leaf tissue remained unchanged for the duration of the experiment (Figure 1B). Many of the isolates (71/124) could spoil both spinach and chicken, 21/124 were only capable of spoiling chicken, 23/124 were only capable of spoiling spinach, and 9/124 were incapable of spoiling either food, even after 7 days incubation (Supplementary Table 2).

Looking at spoilage capabilities at the species level, 100% of *P. aeruginosa* (14 isolates)*, P. furukawaii* (3 isolates) and *P. trivialis* (1 isolates) were capable of spoiling both chicken and spinach (Figure 1C-D). For *P. fluorescens,* 39/54 isolates could spoil both chicken and spinach; 12/54 isolates were only capable of spoiling chicken; 2/54 isolates were capable of only spoiling spinach; and 1/54 was incapable of spoiling either food (Figure 1C-D). For *P. fragi,* 6/20 isolates were capable of spoiling both foods; 8/20 were only capable of spoiling chicken; 3/20 were only capable of spoiling spinach; and 3/20 were incapable of spoiling either food (Figure 1C-D). Of the 23 *P. putida* isolates, only 4 were capable of spoiling both chicken and spinach; 1/23 could only spoil chicken, whereas 14/23 could only spoil spinach; and 4/23 could not spoil either food (Figure 1C-D). For *P. koreensis,* 4/5 isolates could spoil both foods, with the remaining isolate only capable of spoiling spinach (Figure 1C-D). Finally, for *P. chlororaphis* and *P. veronii* all isolates (1 and 2, respectively) were only capable of spoiling spinach and the single *P. alkylphenolica* isolate was incapable of spoiling either chicken or spinach (Figure 1C-D).

We investigated whether the fresh food that the *Pseudomonas* species was isolated from (i.e. meat, seafood or leafy greens; see Supplementary Table 1) had an impact on their ability to spoil chicken or spinach. For isolates from meat products the majority were capable of spoiling chicken (85% of isolates) and spinach (82% of isolates) (Figure 1E-F); for isolates from seafood more were capable of spoiling chicken (95%) than spinach (55%); and for isolates from leafy greens, they were overall much better at spoiling spinach (78%) than chicken (45%) (Figure 1E-F).

Next, we wanted to visualise where live *Pseudomonas* species were located on chicken and spinach during spoilage. To visualise live bacteria, we used the redox-based dyes, 5-Cyano-2,3-Ditolyl-Tetrazolium Chloride (CTC) for chicken and 2,3,5-Triphenyl-tetrazolium chloride (TTC) for spinach. Respiring cells reduce the tetrazolium dye to a formazan crystal, which is visualised in the red fluorescent channel (CTC) or as red visible light (TTC). The chicken samples were also stained with 4′,6-diamidino-2-phenylindole (DAPI) to stain the chicken tissue, then fixed and imaged using wide-field deconvolution fluorescence microscopy. For spinach we used light microscopy and a colour camera to visualise red TTC staining on the green leaf. Since both chicken and spinach samples were washed multiple times prior to imaging, we were only visualising biofilm structures that were strongly attached to the food surface; any loosely attached cells/aggregates and/or slime was removed and not visualised. This allowed us to visualise how the bacteria were interacting with and potentially modifying the tissue structure during spoilage.

We saw destruction of chicken tissue in cases where chicken spoilage had occurred compared to the uninoculated control (Figure 2). This destruction is clearly visible in the DAPI channel images where we observed large voids of varying sizes that extended down into the tissue (Figure 2B-D). Remarkably, most of these voids were filled with CTC-stained *Pseudomonas* biofilms, demonstrating that chicken spoilage is associated with the formation of dense, pillar-like biofilm structures that extend into the chicken tissue (Figure 2C-D). Since DAPI was added to samples pre-fixing i.e. cells were live, the DAPI stain did not penetrate the bacterial cells (Figure 2). Furthermore, the DAPI stain was not localised to nuclei in the chicken tissue, as would be expected in live, intact muscle (Figure 2). This diffuse DAPI staining was also observed in the uninoculated sterile control, however in this sample no voids in the DAPI staining were observed (Figure 2A).

**Figure 2.**
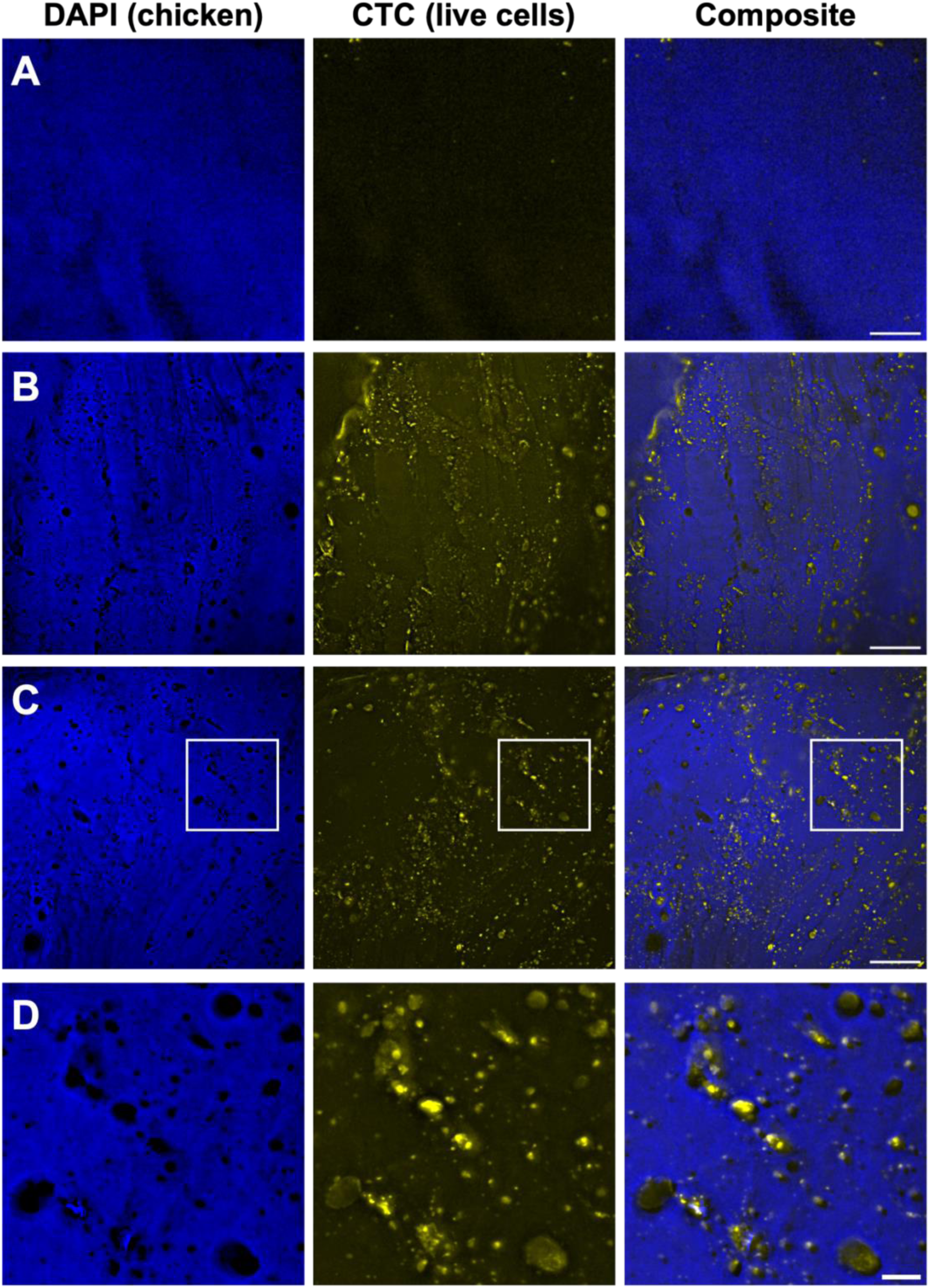
Tissue destruction is associated with biofilm formation on chicken. *In situ* biofilms formed by *Pseudomonas* isolates that are capable of medium-high levels of chicken spoilage were examined using fluorescence microscopy. Voids in DAPI staining were apparent in all samples that had been inoculated with test strains or retained the native microbiota. Most of the voids were filled with biofilm. (A) Sterilised chicken that was uninoculated; (B) unsterilised retail chicken with the native microbial community; (C) sterilised chicken inoculated with *P. fluorescens* isolate #55 (high level spoiler) with full field of view or (D) zoom-in region for each channel (white boxes in C). All samples had been incubated for 2 days at ∼22 °C and then 5-Cyano-2,3-Ditolyl-Tetrazolium Chloride (CTC) was added for 1 h to stain for live bacteria (yellow) and 4’,6-diamidino-2-phenylindole (DAPI) to stain the chicken tissue (blue) and the sample was fixed with paraformaldehyde. Images were obtained using a DV Elite with widefield deconvolution. Images are representative of at least 10 fields of view from two biological replicates. The biofilms formed by other *Pseudomonas* isolates appeared similar to those in B-D. For A-C scale bar = 500 µm, for D scale bar = 100 µm.

To determine if the biofilms formed by *Pseudomonas* isolates on sterilised chicken were representative of the spoilage biofilms formed by microbes present on retail chicken, we allowed fresh chicken to spoil under the same incubation conditions. Macroscopically the samples appeared similar to chicken spoilage in our single-species assays (Figure 1A). Additionally, when we stained these retail chicken samples with CTC and DAPI we observed similar voids in the chicken tissue that were filled with biofilm structures that penetrated the tissue (Figure 2B). This demonstrates that our chicken spoilage assay is a good model for understanding the development of biofilms associated with chicken spoilage.

We also followed biofilm development by *Pseudomonas* isolate #95 that was capable of a high degree of chicken spoilage (Supplementary Table 2). This isolate was inoculated onto sterilised chicken and stained and imaged at days 2 and 4. We used DAPI and CTC stains as above and included the cell impermeant DNA stain, ethidium homodimer-III (EthHD-III), since extracellular DNA (eDNA) is a common component of *Pseudomonas* species biofilms (Okshevsky et al., 2015) (Figure 3). As for Figure 2, there was no alteration in the chicken tissue for the uninoculated sample (Figure 3A), whereas *Pseudomonas* isolate #95 caused significant destruction of the chicken tissue at day 2, which was even more pronounced at day 4 (Figure 3B-C). The biofilms extended in pillar-like structures into the chicken tissue at day 2 (Figure 3B). These structures were also observed at day 4, however, at this time point there were also some large regions (∼100-200 µm in diameter) of tissue destruction which were filled with biofilms (Figure 3C). There was also eDNA and/or dead bacterial cells observed in the inoculated samples that was associated with the pillar-like biofilm structures (Figure 3B-C).

**Figure 3.**
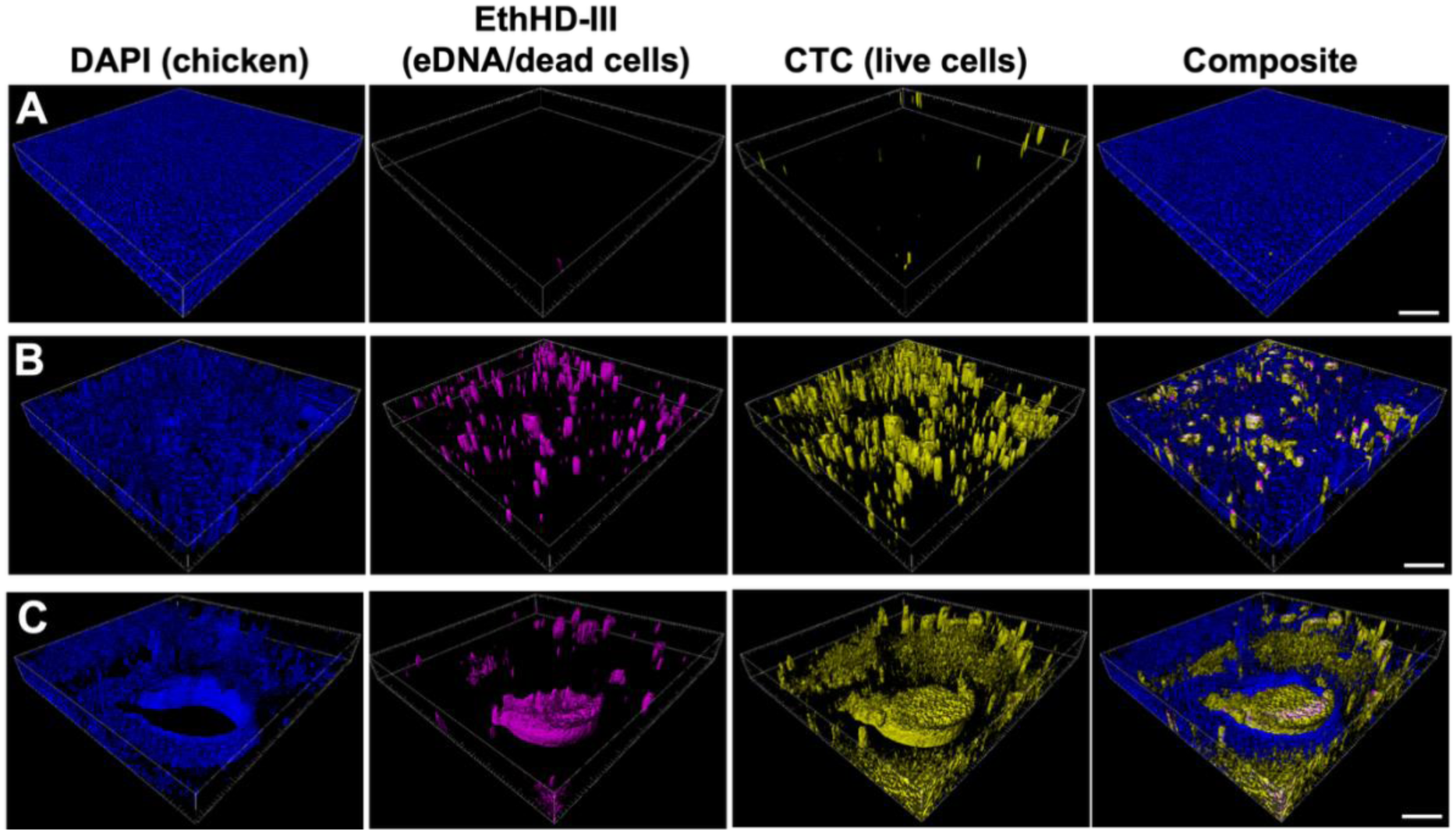
Spoilage biofilm development on chicken. (A) Uninoculated sterilised chicken (day 2 and day 4 images were indistinguishable); sterilised chicken inoculated with *P. aeruginosa* isolate #95 and incubated for (B) 2 days or (C) 4 days. At each time point, samples were stained with 5-Cyano-2,3-Ditolyl-Tetrazolium Chloride (CTC) to stain for live bacteria (yellow), 4’,6-diamidino-2-phenylindole (DAPI) to stain the chicken tissue (blue) and ethidium homodimer-III (EthHD-III) to stain for extracellular DNA (eDNA)/dead cells (magenta) and the sample was fixed with paraformaldehyde. Images were obtained using a DV Elite with widefield deconvolution. The top of each 3D image is the external surface of the chicken, and the bottom side of the image is inside the chicken tissue. Images are representative of at least five fields of view from two biological replicates. Scale bar = 50 µm.

For spinach spoilage we saw clear destruction of the leaf tissue that was often associated with regions containing live bacteria (i.e. TTC stained) that were arranged in biofilm aggregates (Figure 4B-F). For strains with low levels of spoilage whilst we saw many biofilm aggregates on the leaf and rib region, there was minimal leaf destruction (i.e. holes) visible (Figure 4B). For strains with moderate spoilage capabilities, we observed biofilm aggregates on the leaf and rib with more regions of tissue degradation being visible (Figure 4C-D). For strains with high spoilage capacity the cuticle and upper epidermis was degraded (Figure 4E). In these samples we observed few biofilm aggregates on the actual leaf surface and more surrounding the rib region, presumably since this was the region with the highest level of structural integrity that remained in the spoiled leaf. The leaf degradation and biofilms formed by *Pseudomonas* isolates in our model (Figure 4B-E) were similar to spoilage caused by the native microbiota on spinach that had not be sterilised (Figure 4F). This confirms that our spinach spoilage assay is a good model for studying biofilms associated with spinach spoilage.

**Figure 4.**
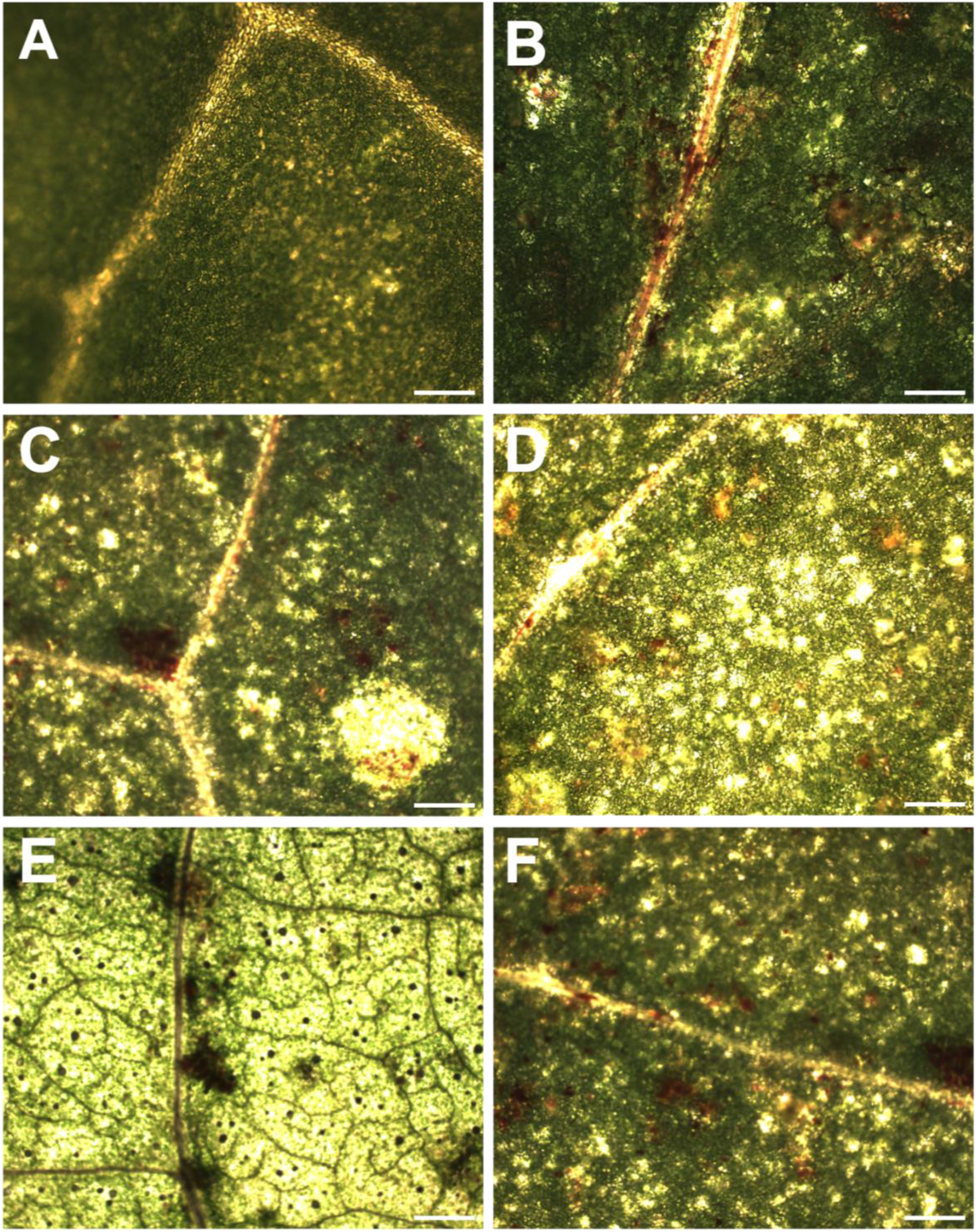
Biofilms formed by *Pseudomonas* species on spinach. Sterilised spinach leaves were inoculated with a single *Pseudomonas* isolate and incubated for 2 days at ∼22 °C. After this time 2,3,5-Triphenyl-tetrazolium chloride (TTC) was added to stain for live bacteria (red). Images were obtained using a colour camera for (A) uninoculated; (B) *P. fragi* strain #11 (low level spoilage); (C) *P. fluorescens* strain #54 (moderate level spoilage); (D) *P. putida* strain #12 (moderate level spoilage); (E) *P. aeruginosa* strain #100 (high level spoilage); or (F) unsterilised spinach with native microbiota (moderate spoilage level). Images are representative of at least five fields of view from two biological replicates. Scale bar = 500 µm.

Our data suggests that spoilage bacteria attach to food surfaces, form biofilms and cause spoilage via tissue degradation. We wanted to further explore the correlations between spoilage and phenotypes associated with biofilm development and secreted enzyme production. It is well established that *in vitro* biofilm formation requires initial attachment to the substratum, which in *Pseudomonas* species is often mediated by twitching and swimming motility (O’Toole and Kolter, 1998). *Pseudomonas* spoilage species secrete several extracellular enzymes, including lipases, proteases and cellulases that are thought to be involved in either tissue degradation and/or food spoilage (Alegbeleye et al., 2022; Barth et al., 2009; Braun et al., 1999). We therefore assayed the panel of *Pseudomonas* isolates for attachment/early biofilm formation, twitching and swimming motilities, and secreted enzyme activities using well-established *in vitro* assays (Supplementary Figure 1; Supplementary Table 3). To determine if *in vitro* phenotypes correlated with spoilage capabilities, we performed separate multiple logistic regression analyses with chicken or spinach spoilage outcome (i.e. spoilage or no spoilage) as the dependent variable and *in vitro* phenotypic data as the independent variables (Table 1). Overall, these analyses show that there is no significant correlation between spinach spoilage and any of the *in vitro* phenotypes (Table 1). There was no significant correlation with any of the *in vitro* phenotypes and chicken spoilage other than secreted protease activity (Table 1).

We also investigated whether the presence of genes encoding secreted enzymes correlated with spoilage capabilities. We looked for the presence of genes encoding known *Pseudomonas* secreted proteases, AprX, AprA, Piv, LasA, LasB and PaaP, secreted lipases TliA, LipC and LipA2 and secreted cellulase BcsZ by performing tBLASTn searches against the whole genome sequences of all isolates using a 70% amino acid identity and 70% coverage cut off (Supplementary Table 4). The genes *aprA, piv, lasA, lasB, paaP* and *lipC* were encoded in 100% of the *P. aeruginosa* isolates, which were all capable of spoiling chicken and spinach. These genes were not found in any of the non-*P. aeruginosa* isolates except for *P. furukawaii* isolates, which possessed *paaP*. To determine if there was a correlation between the presence of genes encoding other secreted proteases, lipases and cellulases (*aprX, tliA, lipA2 and bcsZ*) in the non-*P. aeruginosa* isolates with spoilage capabilities we performed Pearson’s correlation coefficient analyses. This revealed that there was no significant positive correlation for any of these genes and spoilage capabilities, except for *aprX* and chicken spoilage (Supplementary Table 5). Interestingly, almost all isolates that possessed *aprX* (54/58 isolates; 93%) were capable of chicken spoilage. Of the isolates that did not encode *aprX*, 46% of these (24/52 isolates) were capable of chicken spoilage, which suggests that there are other secreted enzymes involved.

## Discussion

It is well established that various members of the *Pseudomonas* genus cause food spoilage and that the relative abundance of these species within the microbial community increases from when food is fresh until it is spoiled (Karanth et al., 2023). Here we wanted to understand how widespread the prevalence of food spoilage capacity is in *Pseudomonas* strains isolated from fresh food. We used a panel of 124 *Pseudomonas* strains and determined their individual capacity to spoil chicken and spinach. We developed models of chicken and spinach spoilage that were excellent representations of the real-world situation as they recapitulated spoilage observed with the native *in situ* communities (Figure 2B, Figure 4F). For the uninoculated controls in both models, we observed no changes in the chicken or spinach tissue over time, which indicates that the observed changes in the spoiled food were due to microbial activities (Figure 1A-B; Figure 2A, Figure 3A, Figure 4A).

To our knowledge this is the most comprehensive study of food spoilage by *Pseudomonas* species. We observed that most isolates were capable of spoiling both foods, with a few isolates displaying specialisation for chicken or spinach spoilage (Figure 1C-D, Supplementary Table 2). Our data also demonstrated that the type of fresh food the strain was isolated from influenced spoilage capabilities: strains isolated from meat were likely to spoil both chicken and spinach, while those isolated from leafy greens were more likely to spoil only spinach (Figure 1E-F).

Interestingly, a small number of isolates were not capable of spoiling either food in isolation (Figure 1C-D, Supplementary Table 2), however since food microbiomes are comprised of mixed communities, it is possible that these isolates may contribute to spoilage as a member of a more complex community.

There has been little exploration of how *Pseudomonas* species interact with food tissues during spoilage. Here we showed that *Pseudomonas* species that were capable of chicken and spinach spoilage formed biofilms associated with significant tissue destruction (Figure 2-4). To our knowledge, our study is the first to visualise *Pseudomonas* spoilage species biofilms on post-harvest spinach. The *Pseudomonas* species biofilms on post-harvest spinach appear similar to previous reports of biofilms formed on (live) leaf surfaces pre-harvest (Morris et al., 1997). As spoilage progresses, there is more degradation of the cuticle and upper epidermis (Figure 1B), and biofilms are found less frequently on the leaf region and are more often localised to the rib region (Figure 4B-F). This suggests that active degradation of the leaf tissue forces the bacteria to either move to another, intact, location on the leaf or rib, or to disperse.

During chicken spoilage, we observed deep penetration of *Pseudomonas* species biofilms into the tissue (Figure 2-3). To our knowledge, this is the first time that spoilage-associated biofilms have been visualised below the surface of the chicken tissue. The ability to penetrate the chicken tissue is likely because the tissue is dead, thus the cell membranes are easily breached and there is no host immune system to protect against microbial invasion. In the current study we are unable to determine if there are specific components of the chicken tissue where cells preferentially attach to then penetrate the tissue. Overall, the biofilms formed on chicken and spinach are unique compared to well-studied *in vitro* biofilms where the substratum is inert and unchanging e.g. on plastic or glass (Sauer et al., 2022). This is consistent with our *in vitro* biofilm assay results, which did not correlate with spoilage capabilities (Table 1). Indeed, previous studies have shown that while *in vitro* biofilm assays facilitate high throughput rapid screening experiments, the conditions and nature of the biofilms formed are typically not a good representation of biofilms formed in real-world settings (Bjarnsholt et al., 2013).

Our results show that chicken spoilage capabilities were only significantly correlated with either *in vitro* secreted protease activity or the presence of a gene encoding a known secreted protease (Table 1; Supplementary Table 5). Of the 92 isolates capable of chicken spoilage, 73 (80%) had either *in vitro* secreted protease activity, or encoded a secreted protease (Supplementary Tables 2-4). However, there remained 19 isolates (20%) capable of chicken spoilage that lacked either *in vitro* protease activity or encoded a known secreted protease (Supplementary Tables 2-4). This suggest that there may be other proteases that are involved in chicken spoilage in these strains. The lack of correlation between other secreted enzyme activity or genes (lipases and cellulase) and spoilage (Table 1; Supplementary Table 5) suggests that these secreted enzymes are unlikely to play an important role in *Pseudomonas* species spoilage of chicken or spinach, or that the environmental conditions in *in vitro* assays are not representative of the *in situ* conditions. It also suggests that there are other secreted enzymes involved in spoilage that target different cellular components. To obtain a detailed understanding of the molecular mechanisms involved in food spoilage, we need to either develop *in vitro* assays and conditions that better mimic the *in situ* conditions and/or study spoilage-relevant phenotypes directly on food surfaces.

## Conclusions

We investigated spoilage capabilities of a large panel of *Pseudomonas* species isolated from fresh food. The majority of these isolates were capable of spoilage of chicken and/or spinach. Spoilage of chicken and leaf tissues was associated with biofilms and significant tissue destruction. To gain a comprehensive understanding of microbial food spoilage requires additional investigations into the roles and interactions between *Pseudomonas* spoilage species and other members of the food microbiome.

## Authorship contribution statement

L.M.M: Writing – original draft; Writing – review and editing; Conceptualisation; Project administration; Supervision; Funding acquisition; Investigation; Formal analysis; Data curation; Visualisation; and Methodology.

J.J.L, R.S., G.J.W, H.A: Writing – review and editing; Investigation; Methodology.

S.J.B and A.E.M: Writing – review and editing; Resources.

C.B.W: Writing – review and editing; Conceptualisation; Project administration; Supervision; Funding acquisition; Investigation; Formal analysis; Visualisation; and Methodology.

## Acknowledgements

The authors would like to thank the Quadram Institute Bioscience Advanced Microscopy Facility (QIBAM) for assistance with microscopy.

## Data availability

Data will be made available on request. *Pseudomonas* genomes from Bloomfield et al. 2024 and from the current study are available in the Sequence Read Archive under Bioprojects: PRJNA973713, PRJNA1248571 and PRJNA1286767.

