## Supplementary Information for "Tissue destruction during food spoilage is associated with the formation of biofilms by *Pseudomonas* species"

**Supplementary Tables**

**Supplementary Table 1.** List of all *Pseudomonas* isolates used in the current study with isolate number, species, the food the strain was isolated from and the corresponding isolate name from Bloomfield et al., (2024) or from this study.

**Supplementary Table 2.** Spoilage scoring for all *Pseudomonas* isolates at 2 and 7 days on chicken and spinach. (-) no spoilage; (+) low spoilage, (++) medium spoilage and (+++) high spoilage.

**Supplementary Table 3.** *In vitro* assay data from biofilm microtitre plate assays, twitching and swimming motility, and secreted protease, lipase and cellulase assays for all *Pseudomonas* isolates.

**Supplementary Table 4.** Presence of genes encoding known *Pseudomonas* secreted proteases, AprX, AprA, Piv, LasA, LasB and PaaP, secreted lipases TliA, LipC and LipA2 and secreted cellulase BcsZ in the isolate panel. These were determined by performing tBLASTn searches against the whole genome sequences of all isolates using a 70% amino acid identity and 70% coverage cut off.

**Supplementary Table 5.** Pearson's correlation coefficient (r) analyses with presence/absence for individual secreted enzyme genes correlated with chicken or spinach spoilage capabilities (from Supplementary Table 2)

### Supplementary Table 5. Pearson's correlation coefficient (r) analyses

**Chicken spoilage capabilities**

| **Gene** | *aprX* | *tliA* | *lipA2* | *Any secreted protease* |
|---|---|---|---|---|
| **r** | 0.5161 | -0.3073 | -0.1072 | 0.5446 |
| **95% CI** | 0.3640 to 0.6414 | -0.4677 to -0.1274 | -0.2887 to 0.08163 | 0.4074 to 0.6577 |
| **R squared** | 0.2663 | 0.09446 | 0.01150 | 0.2966 |
| **P (two-tailed)** | **<0.0001** | **0.0011** | 0.2648 | **<0.0001** |
| **n** | 110[a] | 110[a] | 110[a] | 124[b] |
| **Overall correlation** | positive | negative | none | positive |

**Spinach spoilage capabilities**

| **Gene** | *aprX* | *tliA* | *lipA2* | *bcsZ* |
|---|---|---|---|---|
| **r** | 0.1561 | -0.06982 | -0.1025 | 0.1025 |
| **95% CI** | -0.03207 to 0.3336 | -0.2537 to 0.1190 | -0.2843 to 0.08636 | -0.08636 to 0.2843 |
| **R squared** | 0.02437 | 0.004875 | 0.01051 | 0.01051 |
| **P (two-tailed)** | 0.1034 | 0.4685 | 0.2864 | 0.2864 |
| **n** | 110[a] | 110[a] | 110[a] | 110[a] |
| **Overall correlation** | none | none | none | none |

[a] All non-*P. aeruginosa* isolates

[b] All isolates including *P. aeruginosa*

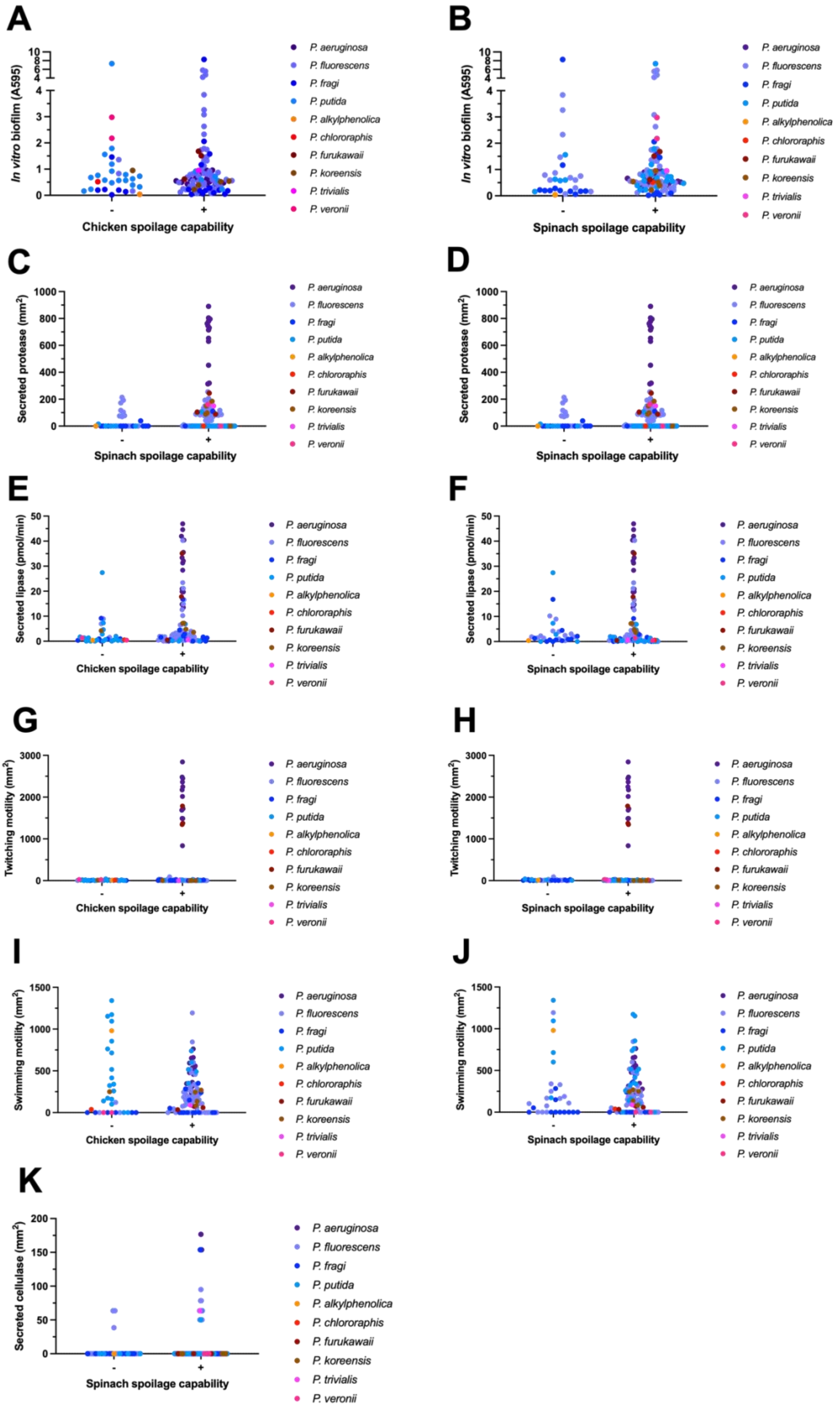
A
In vitro biofilm (A595)
Chicken spoilage capability
B
In vitro biofilm (A595)
Spinach spoilage capability
C
Secreted protease (mm2)
Spinach spoilage capability
D
Secreted protease (mm2)
Spinach spoilage capability
E
Secreted lipase (pmol/min)
Chicken spoilage capability
F
Secreted lipase (pmol/min)
Spinach spoilage capability
G
Twitching motility (mm2)
Chicken spoilage capability
H
Twitching motility (mm2)
Spinach spoilage capability
I
Swimming motility (mm2)
Chicken spoilage capability
J
Swimming motility (mm2)
Spinach spoilage capability
K
Secreted cellulase (mm2)
Spinach spoilage capability
-
+
P. aeruginosa
P. fluorescens
P. fragi
P. putida
P. alkylphenolica
P. chlororaphis
P. furukawaii
P. koreensis
P. trivialis
P. veronii

**Supplementary Figure 1. *In vitro* biofilm, motility and secreted enzyme assays with *Pseudomonas* isolates.** (A-B) *In vitro* biofilm assay data; (C-D) secreted protease assay data; (E-F) secreted lipase assay data; (G-H) twitching motility assay data; (I-J) swimming motility assay data; and (K) secreted cellulase assay data for all isolates grouped by spoilage capabilities for chicken or spinach. Full data is in Supplementary Table 3 and multiple logistic regression analyses on the data are presented in Table 1.
